# An Automated Addressable Microfluidics Device for Minimally Disruptive Manipulation of Cells and Fluids within Living Cultures

**DOI:** 10.1101/688424

**Authors:** Anh Tong, Quang Long Pham, Vatsal Shah, Akshay Naik, Paul Abatemarco, Roman Voronov

**Author notes:** Address correspondence to Prof. Roman S. Voronov, Otto H. York Department of Chemical and Materials Engineering, New Jersey Institute of Technology, Newark, NJ 07102, USA. Electronic mail.

## Abstract

According to the U.S. Department of Health & Human Services, nearly 115,000 people in the U.S needed a lifesaving organ transplant in 2018, while only ∼10% of them have received it. Yet, almost no artificial FDA-approved products are commercially available today – three decades after the inception of tissue engineering. It is hypothesized here that the major bottlenecks restricting its progress stem from lack of access to the inner pore space of the scaffolds. Specifically, the inability to deliver nutrients to, and clear waste from, the center of the scaffolds limits the size of the products that can be cultured. Likewise, the inability to monitor, and control, the cells after seeding them into the scaffold results in nonviable tissue, with an unacceptable product variability. To resolve these bottlenecks, we present a prototype addressable microfluidics device capable of minimally disruptive fluid and cell manipulations within living cultures. As proof-of-concept, we demonstrate its ability to perform *additive* manufacturing by seeding cells in spatial patterns (including co-culturing multiple cell types); and *subtractive* manufacturing by removing surface adherent cells via focused flow of trypsin. Additionally, we show that the device can sample fluids and perform cell “biopsies” (which can be subsequently sent for ex-situ analysis), from any location within its Culture Chamber. Finally, the on-chip plumbing is completely automated using external electronics. This opens the possibility to perform long-term computer-driven tissue engineering experiments, where the cell behavior is modulated in response to the minimally disruptive observations (e.g. fluid sampling and cell biopsies) throughout the entire duration of the cultures. It is expected that the proof-of-concept technology will eventually be scaled up to 3D addressable microfluidic scaffolds, capable of overcoming the limitations bottlenecking the transition of tissue engineering technologies to the clinical setting.

## I. INTRODUCTION

According to the U.S. Department of Health & Human Services, nearly 115,000 people in the U.S needed a lifesaving organ transplant in 2018, while only ∼10% of them have received it. Yet, almost no FDA-approved^1^ artificial organs are commercially available today – three decades after the inception^2^ of tissue engineering and after billions of dollars invested into its development. Therefore, a new approach to biomanufacturing is needed. However, there are major obstacles restricting the progress of complex organ and tissue recreation *in vitro*^3^:

1. Product Size Limitations – due to the lack of an active vasculature and blood circulation within lab-grown tissues, it is difficult to deliver nutrients to / clear metabolic waste from the inner pore space of large (organ-sized) scaffolds. As a result, cell survival in the deep portions of the large scaffolds is compromised due to hypoxia and insufficient nutrient availability. Specifically, scaffolds with dimensions larger than 3.8 x 3.8 x 3.8 mm are classified as *large-volume*, and constructs larger than that (e.g., 5.6 x 5.6 x 5.6 mm) have been found to result in areas of hypoxia after 3 days of culture.^4, 5^ Consequently, out of the three FDA-approved cellular therapies, one (LAVIV injectable fibroblasts for wrinkle treatment) is a suspension of disconnected cells and two (MACI knee cartilage implant and GINTUIT topical treatment of dental wounds) are flat strips of tissue.^1^ No artificial 3D solid organs or complex tissues are currently available.
2. Sacrificial Analysis – due to the inability to sample cells and fluids from within scaffolds nondestructively, and because live long-term 3D microscopy is challenging. This makes it necessary to perform destructive testing at the conclusion of each experiment, such as histological sectioning or crushing the scaffold for plate reader assays. As a result, a different sample must be cultured for each new time point. This balloons the cost of experiments and slows down the scientific progress tremendously.
3. Product Variability – due to the absence of a native supervision over cell behavior and lack of access to them post seeding, there is no orchestration over their actions within the scaffolds. For example, even if one were to bioprint (i.e., deposit the cells into precise locations within a scaffold) the perfect artificial tissue, the cells within it would be free to do a number of undesirable things afterwards: a) migrate away uncontrollably,^6^ b) differentiate into the wrong tissue type (e.g., a patient grew mucus tissue in her spine, as a result of stem cell therapy),^7^ and c) deposit tissue in the wrong locations and occlude the scaffold pores. All this leads to nonviable tissue and poor product consistency. In fact, a survey of 16 big pharmaceutical companies found that “*product consistency is possibly the single greatest challenge facing the field of regenerative medicine*”.^8^
4. Technology Adoption Barriers – Even if one could generate the perfect tissue in a laboratory, training hospital staff in custom culturing protocols for each new product remains another critical hurdle holding back biomanufacturing technologies from entering the market.

In this study, we hypothesize that in order to overcome these obstacles, an ideal scaffold should be composed of the following elements: (1) Active “Vasculature” for distributing metabolites and clearing waste throughout *organ-sized* scaffolds; (2) Nondestructive Sampling of the cells and of the fluids within them for *ex-situ* chemical analysis; (3) Long-term Live Microscopy observation of the cell behavior and tissue growth; (4) Tissue Modulation via cell and bio-active chemical (e.g., chemo-attractants, growth and differentiation factors, drugs, etc.) delivery, in order to minimize product variability; and (5) Automated Spatiotemporal Control over the tissue development in a closed-loop manner, based on optical and chemical assaying feedback, in order to enable computer-driven culturing. The overall idea is depicted in Figure 1.

**Figure 1.**
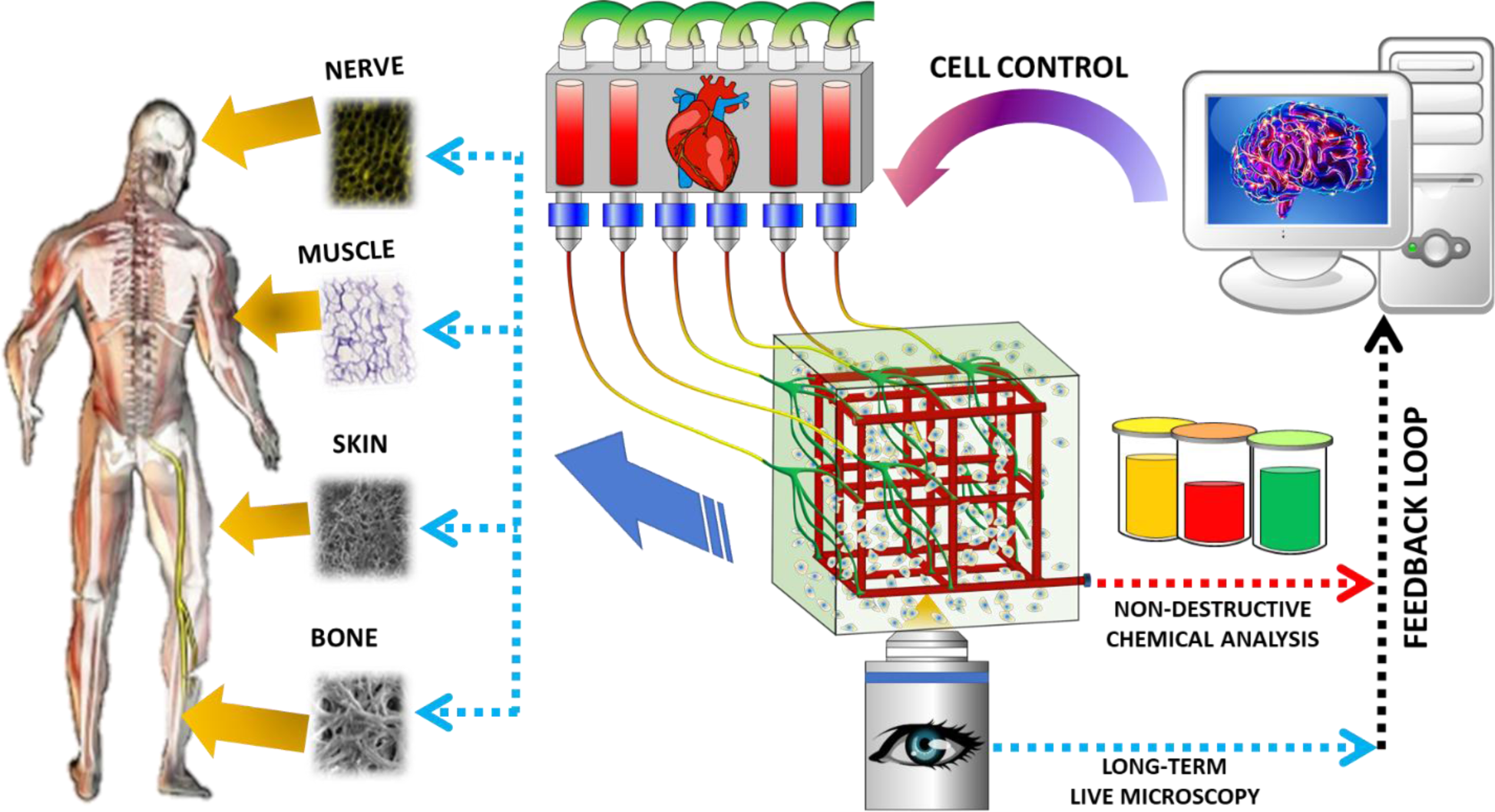
Hypothesized ideal scaffold with active “vasculature” that enables targeted real-time fluid and cell manipulation within the cultured tissue. External pumping acts like a “heart”, while the whole process is orchestrated by a computer acting like a “brain”. The computer’s closed-loop responses are based on feedback from non-destructive chemical analysis and long-term live microscopy throughout the entire duration of the culture. Viable artificial tissue is cultured automatically and reproducibly.

We further hypothesize that these goals can be achieved by merging microfluidic and scaffold technologies. Interestingly, microfluidics technologies share many characteristics with conventional biomanufacturing techniques, while at the same time lacking their major bottlenecks (see Table 1). Namely, they can seed cells with precision, perform nondestructive localized chemical sampling and are transparent to microscopic observation. Moreover, the microfluidic substrates can be fabricated from a wider range of materials, and with even more precision than their bioprinted counterparts, because there is *no danger of damaging the cells* during the fabrication process (since they can be *flowed in afterwards*). Most importantly, the active “vasculature” of micro-channels allows continuous nutrient delivery / waste removal and enables targeted modulation of cell behavior in a closed-loop manner.

**Table 1.** Comparison of existing biomanufacturing technologies (rows 1-3) with the proposed microfluidics approach (bottom row). Green = key advantages, Red = fundamental limitations; Yellow = neutral advantage.

| Tech Type | Material | Geometry | Seeding | Viability | Vasculature | Analysis | Controls |
| --- | --- | --- | --- | --- | --- | --- | --- |
| Artificial Scaffold | Any | Poor to Precise | Random | Excellent | Passive | Bulk | Bulk |
| De-cellularized Organ | Natural | Natural | Random | Excellent | Passive | Bulk | Bulk |
| Bio-printing | Very-Limited | Precise | Precise | Poor to Good | Passive | Bulk | Bulk |
| Microfluidics | Less-Limited | Precise | Precise | Excellent | Active | Localized | Localized |

In contrast, bioprinting is limited to materials that are compatible with cells. The materials are typically soft (e.g., hydrogels) so that they can be extruded, and later hardened via crosslinking. Furthermore, the extrusion can worsen the cell viability, because the cells must survive the manufacturing process that may involve high shear stresses, thermal stress and/or toxic crosslinking agents. Moreover, the manufacturing resolution of the bioprinters is limited by the fact that the nozzle must be big enough to allow the cells to pass through it seamlessly (i.e., typically ∼100 m diameter). Likewise, de-cellularized organs are limited to just the natural extra-cellular matrix material and structure. And for both, this and the artificial scaffold, methods there is little control over where the cells are seeded. Finally, all three of the conventional approaches to tissue engineering have *passive* pores, which cannot be used for in-situ cell analysis or control.

Therefore, the microfluidics scaffolds provide numerous advantages over the conventional tissue engineering approaches. Furthermore, although fabricating microfluidics devices has been a major bottleneck to their adoption by the industry, 3D printers are on the verge of being able to manufacture them.^9, 10^ Hence, it makes sense to integrate scaffolds with the microfluidic technologies, in anticipation of the near future when 3D printing will facilitate their translation to the market.

To that end, several such attempts have been made in the past.^11–18^ However, these mostly focused on developing the *materials* and the *fabrication techniques* for manufacturing of the microfluidic scaffolds, while the plumbing necessary for the targeted fluid and cell manipulation within them has not been designed. Specifically, the internal plumbing of such scaffolds should contain dedicated ***ports***, distributed at targeted locations (from here on termed as “addresses”), in order to enable the localized nondestructive manipulation (i.e., delivery/probing) of cells and fluids within the cultures. Yet, this is a significant challenge because the plumbing should be both *scalable* to *large* (organ-sized) scaffold sizes and at the same time be capable of maintaining a *high density* targeting throughout its pore space. The problem is illustrated using a 2D example in Figure 2A.

**Figure 2.**
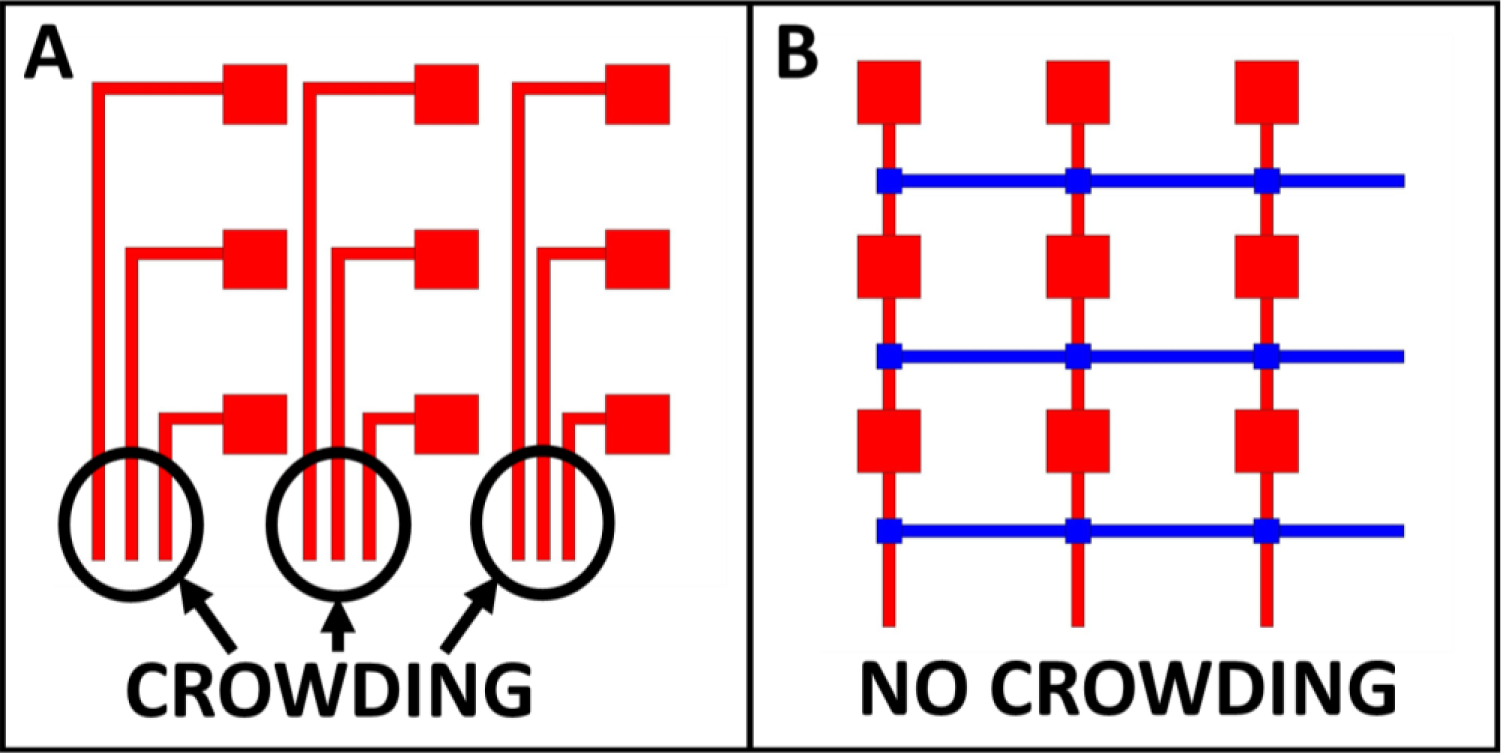
Difference in scalability and resolution of addressable microfluidics arrays, illustrated using a 3 x 3 grid example: **(A)** Inefficient X*Y scaling, and resolution limitations due to crowding of the supply channels; **(B)** Efficient (X+Y) scaling, and resolution is not limited by the crowding. Red = flow channels through which cells or chemicals are delivered and/or sampled; Blue = blocking (i.e., valve actuating) channels

In this figure, a *naive* approach is shown, where a 3×3 grid of microfluidic ports is actuated via a separate channel dedicated to each of the addresses. This suffers from: a) Poor scaling – the number of flow channels required to actuate each individual address in the grid scales as X*Y, which is the worst case scenario; and b) Crowding - the spacing available for the channels is limited by the separation distance between the neighboring columns of addresses (which should be as small as possible, in order to ideally be able to manipulate and analyze the culture with a single cell spatial resolution). Fortunately, an old concept in the microfluidics field solves this problem by including orthogonally-blocking channels,^19–21^ shown in blue in Figure 2B. By using this combination of flow and blocking (i.e., valve actuating) channels, (X+Y) scaling is achieved (i.e., only 3 flow + 3 blocking channels are required, as opposed to a total of 9 dedicated flow channels in the X*Y scaling); and the resolution is no longer limited by the crowding.

Thus, the “addressable” microfluidic technologies, such as in Figure 2B, yield the *best possible scaling* (e.g., microfluidics chips with over 1000 addresses have been made ^22–25^), and the address density is *no longer limited by the crowding*. Therefore, this type of plumbing is the best choice for organ-sized tissue cultures. Yet, it has not been used in microfluidic scaffolds before. Instead, the addresses typically serve as separate *chambers* on chips used for multiple parallel experiments: for example, an array of cell culturing chambers for high-throughput drug testing,^19^ a droplet-based device for multi-parameter analysis of single microbes and microbial communities,^20^ and a stencil for protein and cell patterning of substrates.^21^

However, here we are interested in addressable cell and fluid manipulation *within a single* culture (as opposed to each address corresponding to separate isolated chambers), which has not been done before. Furthermore, we want the device to be *automated*, in order to enable computers to perform the manipulations over long-time culturing experiments. To that end, the remainder of the manuscript presents our proof-of-concept platform, which adapts the addressable microfluidic plumbing and automation, in order to perform the minimally disruptive cell and chemical manipulations needed to revolutionize the tissue engineering scaffolds.

## II. METHODS AND EXPERIMENT

### II.1 Master Mold Fabrication

The device was fabricated via a multilayer soft-lithography technique^26^ using a custom-built UV mask aligner.^27^ The device’s design is discussed in detail in Section III.1 (see Figure 6 and Figure 7). The molds for the device was fabricated using negative photoresist (SU-8, Microchem, MA) and positive photoresist (AZ^®^ P4620, Integrated Micro Materials, TX). First, the microscale pattern was sketched using AutoCAD (Autodesk, Mill Valey, CA) and printed at 50,800 dpi on a transparency (Fineline Imaging, Colorado Springs, CO) to generate a high-resolution photomask. Initially, the 4-in silicon wafers (University Wafer, Boston, MA) were washed carefully with diluted soap, rinsed with Acetone; Methanol; and DI-water (AMD solvents), dehydrated at 180 °C for 15 minutes. Subsequently, the wafers were cooled down to room temperature and treated with Hexamethyldisilazane (HMDS) (VWR, Radnor, PA) to enhance the photoresist adhesion. The procedures to create the master mold for each layer of the microfluidic device are describe below:

#### Payload Layer

The master mold for the Flow Layer consists of two types of photoresists, positive photoresist AZ^®^ P4620 for the flow channels and negative photoresist SU-8 2150 (Microchem, MA)for the addressable ports. **Round profile flow channels** (See Figure 3A) **-** AZ^®^ P4620 was spin coated at 1400 rpm, soft-baked at 90 °C for 10 minutes. Then the second layer of AZ^®^ P4620 was spin coated on the same wafer to reach the target coated layer of 28-μm, soft-backed at 90 °C for 1 hour. Subsequently, the wafer was rehydrated overnight inside the oven at 37 °C with an opened water tray (12 hours), exposed to UV light (exposure dose: 2800 mJ/cm^2^), and developed by immersing them in a vessel containing AZ® 400K developer for 10 to 15 minutes (Integrated Micro Materials, TX) to form 28-μm height features. The round profile of the channels was created by baking the wafer with AZ^®^ P4620 features at 150 °C on a programmable hot plate for 15 hours, starting at 65 °C with heating ramp rate of 4 °C/h. **Addressable Ports** (See Figure 3B) **-** SU-8 2150 was spin-coated directly on the same wafer at 1250 rpm, aligned with the first pattern using the custom mask aligner,^27^ exposed to the UV (exposure dose: 600 mJ/cm^2^), and by immersing them in a vessel containing SU-8 developer for 30 minutes to generate 550 μm-height square features for the addressable ports. The developed photoresist was fully crosslinked at 180 °C for 2 hours, cooled down to room temperature, and treated with Perfluorodecyltrichlorosilane (FDTS) (Alfa Aesar, MA) inside a vacuum desiccator chamber for 4 hours.

**Figure 3.**
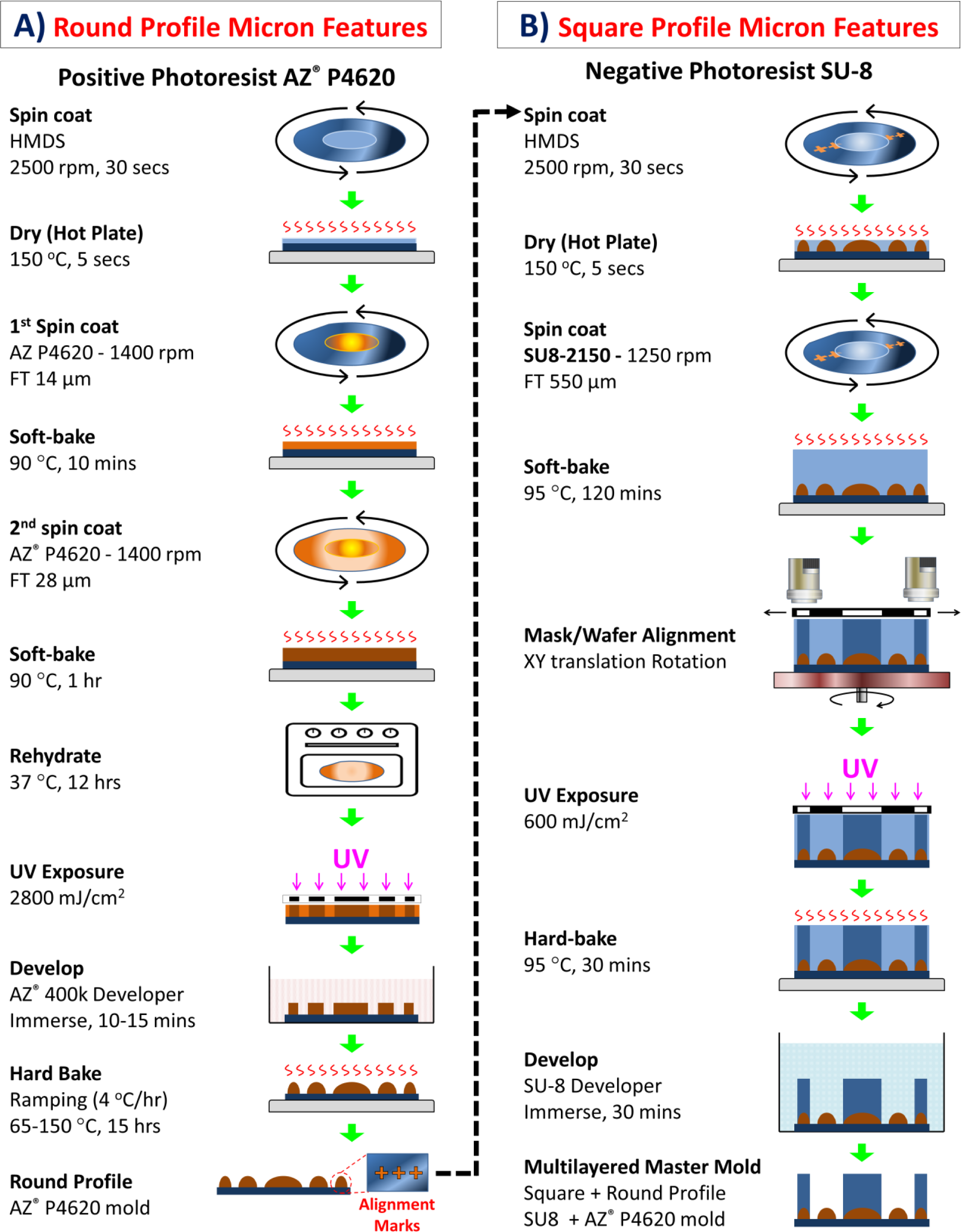
Step-by-step fabrication process of a multilayered master mold, which composed of two different heights/profiles (round and square) of micron-sized features on a 4-in silicon wafer. Basically, the fabrication follows a standard photolithography technique involves spin coating, UV exposing, and developing. **(A)** The positive photoresist AZ^®^ P4620 template was first created and followed by reflowing the developed photoresist by baking the at 150 °C to achieve the round profile features. The alignment marks were also imprinted on the two sides of the wafer, which will be used in the next step. **(B)** SU-8 photoresist layer is patterned on the same wafer. Prior to UV exposure, the alignment marks on template photomask and on the silicon wafer are aligned using the custom mask aligner.^27^ *Abbreviations: FT = Film Thickness*.

#### Control Layer

There were two Control Layers in the addressable device. The first one was used to actuate the valves in the Payload Layer (Figure 4 and Figure 6A), and the second one was to open/close inlets and outlets of the Culture Chamber (Figure 4 and Figure 7). The photomasks were switched respectively for each layer and the procedure to create the master mold for the Control Layer remained the same. Specifically, SU-8 2050 (Microchem, MA) was spin-coated at 1500 rpm, exposed to UV light (exposure dose: 240 mJ/cm^2^), and developed on a 4-in silicon wafer to generate a 120-µm height pattern. The developed photoresist was fully crosslinked at 180 °C for 2 hours, and then slowly cooled down to room temperature.

**Figure 4.**
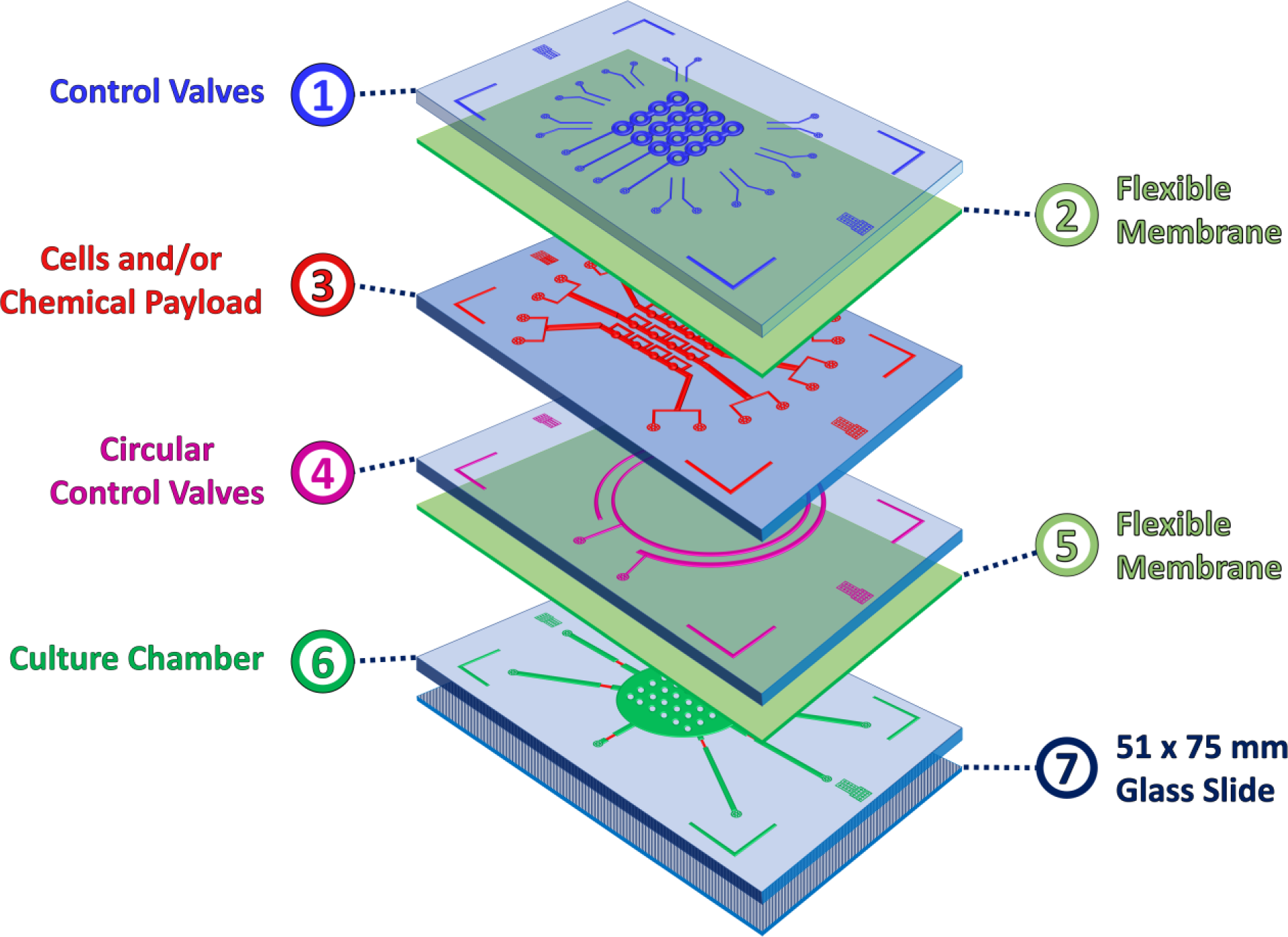
The Multilayered Addressable Microfluidic Device consists of 7 layers. **(1)** Cells and/or Chemical Payload’s Control Layer; **(2)** 35 μm Flexible Membrane; **(3)** Cells and/or Chemical Payload with Addressable Ports Layer; **(4)** The Control Layer for the Culture Chamber; **(5)** 35 μm Flexible Membrane; **(6)** Culture Chamber; and **(7)** 51 x 75 mm Glass Slide.

#### Culture Chamber

The master mold for the Culture Chamber also consisted of two types of photoresists, positive photoresist AZ^®^ P4620 for features, which overlapped with the control Valve Layer, and negative photoresist SU-8 2150 for the cell Culture Chamber and non-overlapping flow channels. The round profile features were created by following the same procedure as the round profile flow channels for the Payload Layer (see above). Then, SU-8 2035 (Microchem, MA) was spin coated at 1500 rpm, exposed to UV light (exposure dose: 210 mJ/cm^2^), and developed on a 4-in silicon wafer to generate 85 µm height square pattern. The developed photoresist was fully crosslinked at 180 °C for 2 hours and then slowly cooled down to room temperature.

### II.2 Microfluidic Device Fabrication

Different Polydimethylsiloxane (PDMS) Sylgard 184 (Midland, MI) layers of the device were generated using soft lithography. The elastomer with a base-to-agent ratio of 10:1 was poured over the photo-patterned mold to reach the thickness of 5 mm, 1 mm, and 2 mm for the payload’s Control Layer, the culture’s Control Layer, and the Culture Chamber, respectively. Then the PDMS casted molds were degassed inside the vacuum desiccator chamber for 2 hours and followed by curing on a hotplate at 65 °C overnight (12 hours). The PDMS flexible membranes (Figure 4 and Figure 6B) of 35 μm-thickness were created by spin-coating the PDMS with 20:1 base-to-agent ratio onto 4-in silicon wafer at 2500 rpm for 60 seconds, and then baked at 65 °C for at least 1 hour. The cells and/or chemical Payload Layer was created by following an established PDMS stencil procedure.^21^ Then all of the layers were peeled off from the master molds, washed with diluted soap, rinsed with AMD solvents, dried on a 180 °C hotplate, treated with air plasma, and bound to each other using our custom-built PDMS desktop aligner^28^ to form the multilayered microfluidic device (Figure 4). The order of binding single layers to form the multilayered microfluidic device was as follows: 1) Cells and/or Chemical Payload’s Control Layer; 2) 35 μm Flexible Membrane; 3) Cells and/or Chemical Payload with Addressable Ports Layer; 4) The Control Layer for the Culture Chamber; 5) 35 μm Flexible Membrane; 6) Culture Chamber; and 7) A biopsy punch (Electron Microscopy Sciences, PA) with a diameter of 0.5 mm was used to create inlet and outlet ports for tubing connections. The whole device was bound onto a 51 x 75 mm glass slide (Corning^®^, NY) using air plasma treatment.

### II.3 Pumping Automation, Experimental Setup, and Microscopy

Here we employ a modification of an open-source programmable pneumatic pumping technology, developed for operation and automated control of single- and multi-layer microfluidic devices.^29–31^ Following this design, we built a custom pneumatic pumping system based on modular industrial automation components made by WAGO (see Figure 5A). Specifically, the core of the setup consisted of an Ethernet-based programmable WAGO-I/O-SYSTEM 750 logic controller, and an 8-channel digital output module (WAGO Kontakttechnik GmbH & Co, Minden, Germany) that allows the controller to drive 24 V Festo (MH1-A-24VDC-N-HC-8V-PR-K01-QM-APBP-CX-DX, Festo, Germany) miniature pneumatic solenoid valves (See Figure 5B). The solenoid valves were connected to a custom DYI pneumatic pumping system (see Figure 5C), which is used to manipulate cells and fluids at the addresses. In order to avoid contact between the solenoid valves and the pumped liquid, the system contains machined reservoirs (see Figure 5D) that prevent water from backing up into the former. The solenoid valves are actuated by sending 24 V signals to them via an eight-channel digital output module (Wago 750-530). The “on” and “off” positions of the solenoid valves correspond to the “open” and “close” states of the valves on the chip. Switching from “open” to “close” means changing the pressure inside the on-chip valve from atmospheric pressure to 20 psi, respectively. The device was connected to PYREX® 100 mL Round Media Storage Bottles (Corning^®^, NY) with Solvent Bottle Cap - GL45 - 5 Luer ports (Cole-Parmer, IL) and house-air via Tygon tubing (Cole-Parmer, IL) of 0.02-in inner diameter. The valves in the Control Layers were also automatically actuated via the miniature pneumatic solenoid valves that were operated by the programmable WAGO controller. The controller was connected to a computer via an Ethernet interface. First, a sequence of patterns to be addressed on the device was specified by a user using a custom Matlab^®^ graphical user interface (GUI) (see Figure 5E). Then these patterns were sent to the WAGO module to toggle the Festo valves, which in turn selectively actuated the on-chip valves at the preset locations (specified by the pattern created above) for the delivery of the payloads and collection of samples through the ports.

**Figure 5.**
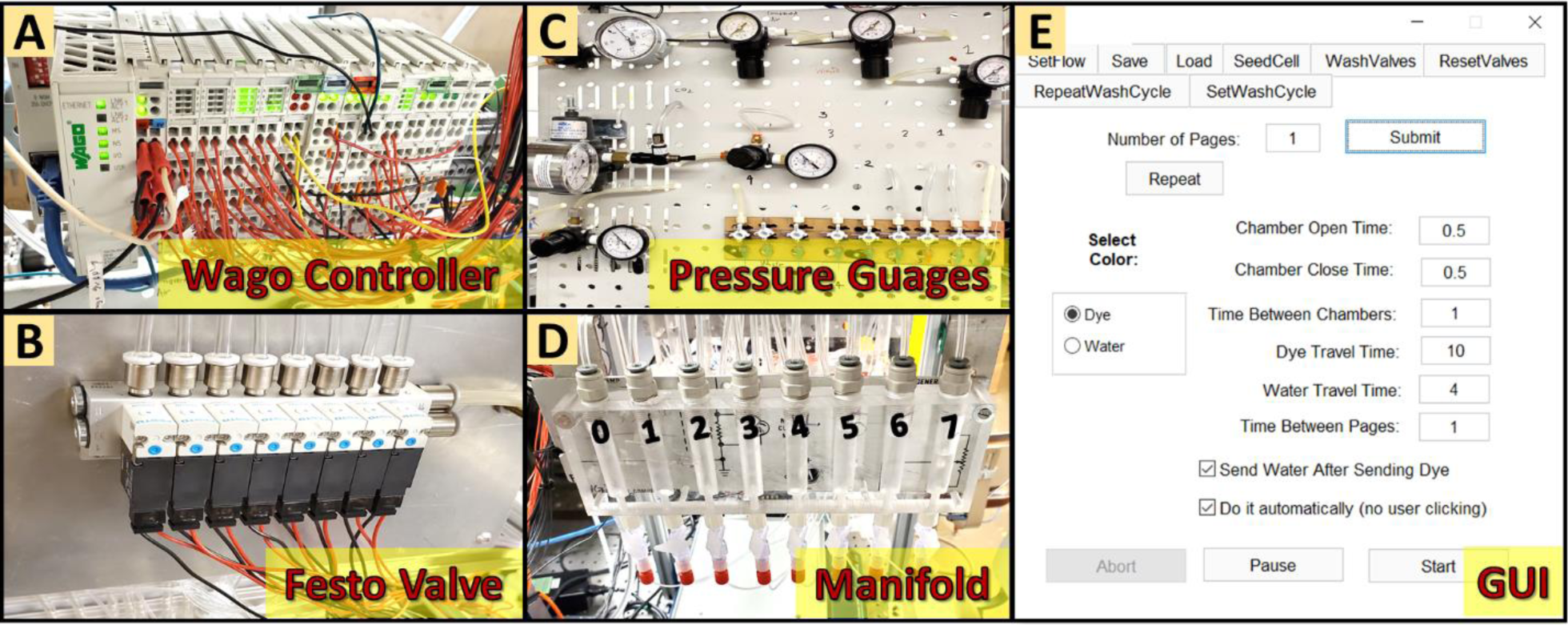
Automated pumping system. **(A)** Wago controller; **(B)** Manifold with 8 Festo solenoid valves; **(C)** Pneumatic pumping system; **(D)** Water storage machined reservoirs for control channels; **(E)** Custom GUI for controlling the solenoid valves via the Wago controller.

The device was mounted on an automated microscope (IX83, Olympus, Japan) equipped with a XY motorized stage (96S106-O3-LE2, Ludl, NY), and a custom temperature control setup. Images were acquired using a digital CMOS camera (Orca Flash V4, Hamamatsu, Japan). The image acquisition was performed using a custom Matlab^®^ (MathWorks Inc., Natick, Massachusetts) GUI.^32^ The image-processing was parallelized using the Matlab^®^’s Distributed Computing Toolbox in order to enable on-line analysis of the culture.

### II.4 Demonstration of *Fluid* Manipulation within the Device

Since many biochemical agonists are colorless / undetectable by regular microscopes, food dye (Assorted Neon!, McCormick, Baltimore, MD) was used in order to create a visual demonstration of the fluid manipulations within the microfluidic device. The dye was added into the media bottles connected to the flow channels of the device via Tygon tubing. Air pressure of ∼ 1 psi was used to drive the fluid into the device and across the flow channels. On-chip valves were connected to a pressure source of ∼ 20 psi in order to fully obstruct the flow when the valves were in a “closed” state. Matlab^®^ code was used to preset the pattern in which the dye was delivered and sampled. For the latter, the one outlet of the payload channel was connected to vials under negative pressure via Tygon tubing, in which the chemicals (food dyes) were withdrawn back via a port at any (either at the same or at a different) location of the 4×4 array via pressure reversal into the sample collecting vials for further analysis. Time-lapse video of the dye delivery was recorded using a compact digital microscope (AD4113T, Dino-lite, Torrance, CA).

### II.5 Demonstration of *Cell* Manipulation within the Device

#### Device Preparation for Cell Culturing

The device was autoclaved at 121 °C for 60 minutes to completely cure the non-crosslinked oligomers inside the bulk PDMS, evaporate the remained solvent from curing agent, and sterilize the PDMS device prior to the adhesive surface treatment and cell seeding. Subsequently, the cell Culture Chamber of the device was coated with fibronectin 10ug/mL and the internal surfaces of the microfluidic channels were treated with 2% Bovine Serum Albumin (BSA) (Sigma, MO) at 25 °C for at least 10 hours inside the UV chamber to maintain the sterile condition of the PDMS device. In this case, 2% BSA solution was used to prevent the adhesion of the cells to the microfluidic channel’s surface.^33, 34^

#### Cell Preparation

Culture media was prepared from Minimum Essential Medium (MEM) (Sigma, MO) supplemented with 10% (v/v) fetal bovine serum (FBS) (VWR, Radnor, PA) and 1% (v/v) penicillin-streptomycin (10,000 U mL-1) (Thermofisher, Waltham, MA). Basal media was composed of MEM supplemented with 1% (v/v) penicillin-streptomycin. For incubation in 5% CO_2_ atmosphere, media was buffered by 26 mM sodium bicarbonate (Sigma, MO). CO2-independent media buffered by 20 mM HEPES (Sigma, MO) was used for microscope stage-top experiments. Mouse Bone Marrow-Derived Mesenchymal Stem Cells (MSCs) (S1502-100, strain: C57BL/6) and Mouse Embryo Fibroblasts (NIH/3T3) (ATCC^®^ CRL-1658TM, Strain: NIH/Swiss) were used for this study. The selected cell type was suspended in a pre-warmed complete growth Minimum Essential Medium (MEM) medium, supplemented with 10% fetal bovine serum and Gentamicin at the concentration of 50 µg/mL, to reach the cell concentration of 5 x 10^6^ cells mL^-1^. Initially, the cells were trypsinized from a T-75 cell culture flask by adding 2 mL of 1x Trypsin/Ethylenediaminetetraacetic Acid (EDTA) (0.25%, 0.2 g L−1 EDTA) for 3 min. MEM culture media (8 mL) was added to neutralize the trypsin/EDTA activity. The cell suspension was centrifuged at 1000 × g for 2 min. The supernatant was removed by aspiration and the cell pellet was re-suspended in 5ml of Cell-tracker fluorescent dye (CellTracker™ CM-DiI (Invitrogen™, MA) in Phosphate-Buffered Saline (PBS) (1X without calcium and magnesium) (Sigma, MO) for: 1) red fluorescent cell membrane labeling (λ_exc_ = 553 nm / λ_em_ = 570 nm); CellTracker™ Green CMFDA Dye (Invitrogen™, MA), or 2) green fluorescent cell cytoplasm labeling (λ_exc_ = 492 nm / λ_em_ = 517 nm). The cells suspension was incubated at 37 °C for 45 minutes and then centrifuged at 1000 × g for 2 min. The supernatant was removed by aspiration and the cell pellet was re-suspended in MEM culture media to reach the desired concentration [5 x 10^6^ cells mL^-1^]. Finally, the cell suspension was ready for the cell seeding procedure.

#### Additive Manufacturing

10 mL of the suspended cells was added to a dispensing bottle with 4 ports that were connected to the cell and/or chemical Payload Layer channels of the device. The bottle was then pressurized via 5% Carbon Dioxide at 6-7 psi, under a constant shaking motion at 210 rpm. The on-chip valves were connected to a pressure source of ∼ 20 psi in order to fully block the flow to an address when the valves are in a “closed” state. The Matlab^®^ GUI was used to predetermine the patterns, in which the cells were seeded via the addressable ports. The fluorescent and time-lapse images were captured using a fully automated Olympus IX83 microscope fitted with a 20X phase-contrast objective (UPLFLN20XPH, Olympus, Japan), a CMOS camera (Orca Flash 4.0 V2, Hamamatsu, Japan). Time-lapse images were automatically captured at a 15 minutes interval for a duration of 30 hours. For each time step, 121 tile images (size: 662.07 x 662.07 µm) were acquired at different locations, stitched, and stabilized using an in-house Matlab^®^ 2016b code (MathWorks, Inc., Natick, MA). During the acquisition, the culture media was automatically refreshed every 5 hours via the Culture Chamber’s flow channels of the device by actuating the control valves in that layer “on” and off”. The composite images of the patterned cell co-cultures created by seeding two different cell types in different locations were assembled using the ImageJ software (National Institutes of Health).^35^

#### Subtractive Manufacturing

The suspended cell solution at a concentration of [0.6 x 10^6^ cells mL^-1^] was delivered into the Culture Chamber until the cells were uniformly seeded. After cultivation at 37 °C for 3 h, PBS (1X without calcium and magnesium) was used to flush the cell Culture Chamber for 30 seconds and trypsin (0.25% Trypsin 0.2 g L^-1^ EDTA) was then delivered into the Culture Chamber through user-specified addresses for 10 minutes. The bond cleavage effectiveness was gauged based on real time observation of cell morphology via microscopy. After the digestion, the lifted cells were flushed away by continually flowing the PBS through the same address. Pulsatile flow with a periodicity of every 0.5 second was used with in order to facilitate the cell detachment from the substrate floor of the Culture Chamber.

#### Minimally-Disruptive Cell Biopsies

The cells were uniformly seeded in the Culture Chamber and were treated with trypsin for 5 minutes (following the procedure in the previous section - **Subtractive Manufacturing**) until they were partially detached. Eventually, the detached cells were collected from the culturing chamber through the microfluidic ports via pressure reversal applied at the outlet of the payload channel. Finally, the cells were stored in a vial, so that they could be sent away for ex-situ analysis.

## III. RESULTS

### III.1 Redesigned Microfluidic Plumbing for Addressable Access to the Culture Chamber

In this manuscript, we have realized an envisioned proof-of-concept automated microfluidic platform, capable of minimally-disruptive XY fluid and cell manipulations within live 2D cultures. To do this, we used a combination of micro-sized flow channels and blocking pneumatic valves, in order to actuate the individual addresses *independently* of each other. After going through multiple iterations of modifying the published addressable plumbing designs,^19–21^ we have converged upon the result shown in Figure 6A. There, the addressable array has a 4×4 size for simplicity, though the actual grid size is a free parameter. Each of the addresses in the array are shown as red-discs, which are surrounded by O-shaped pneumatic valves (shown as blue circles). When the valve is “*closed”* (see the inset in Figure 6A), the fluid traveling through the red channels is re-routed around the address via a thin bypass channel (also labeled in red). However, when a valve is “*open”*, the corresponding address can either deliver or withdraw (depending on the direction of the flow in the red channels) the fluid carrying a chemical payload, and/or cells, to/from the Culture Chamber below (which is shown at #4 in Figure 6B).

**Figure 6.**
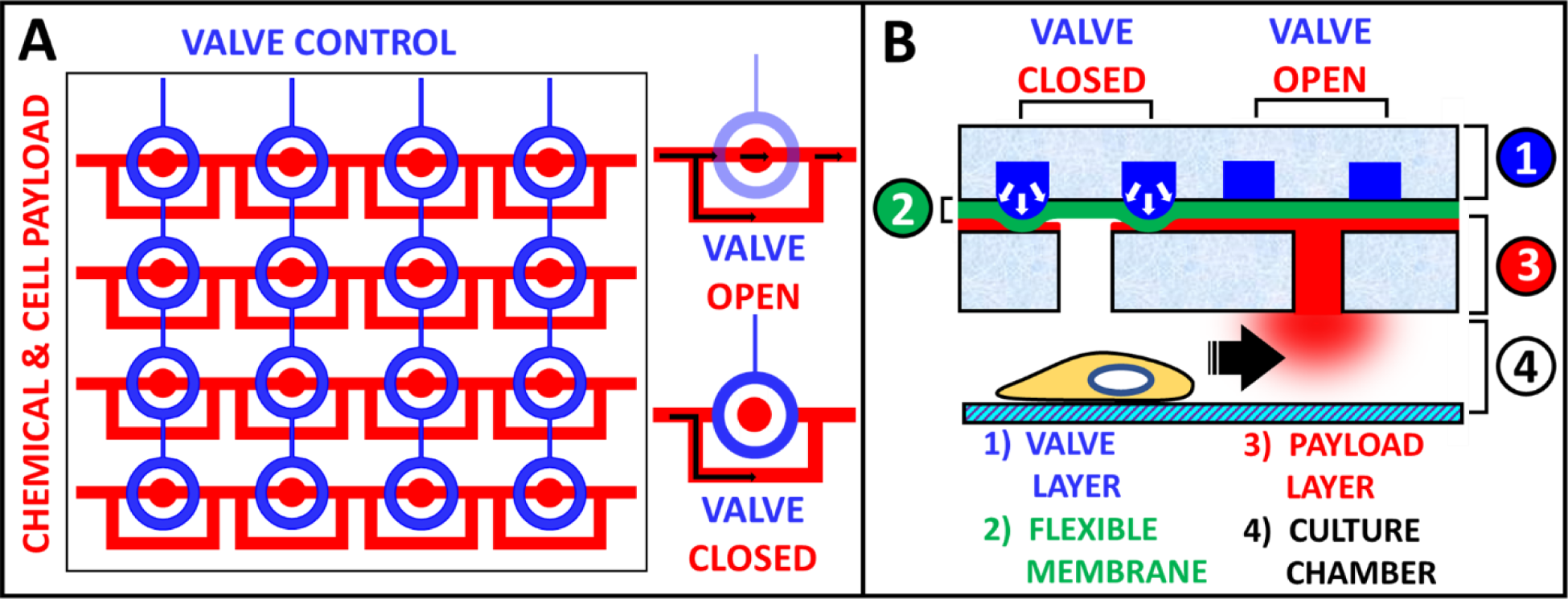
S*hematic of the Addressable Microfluidic Plumbing developed in this manuscript. **(A)** Top XY view of a 4×4 matrix of the “addressable” ports. Flow channels are shown in red and blocking valve channels are in blue. Inset shows that a port is active when the valve is “open” and bypassed when it is “closed”. Black arrows indicate the direction of flow, which assumes a payload* delivery to *(as opposed to* sampling at*) an address. **(B)** Z cross-section of the same device, showing how a cell neighboring an active microfluidic port is attracted towards it due to the chemoattractant released into the Culture Chamber at that location. At the same time, it is shown that the inactive port directly above the cell does not affect its behavior, since no chemical payload is being delivered through this address*.

Figure 6B is a Z cross-section of the device, which shows that it consists of 4 main layers (from top to bottom): a Valve Layer, a thin Flexible Membrane, a Flow Layer, and a cell Culture Chamber. The action of the O-shaped valve is also shown in the same figure: in the *“closed”* state, the pressurized valve expands, causing the flexible membrane to block flow to the address; conversely, in the “*open*” state, the flow is allowed to enter the address freely, where a microfluidic port then connects it with the Culture Chamber below. As an example, a chemical payload can be delivered through this port in order to attract a neighboring cell. At the same time, the *closed* port directly above the cell would not affect its behavior since no chemical is being released at that location. Furthermore, the same ports can be used to sample the culture media and seed or collect cells at different locations throughout the Culture Chamber.

Additionally, the Culture Chamber is supplemented with side channels for replenishing the media, uniform seeding of the cells and flushing of the chamber. Figure 7 shows a zoomed out top view of the Culture Chamber (labeled in red) and two surrounding circular valves (labeled in blue) that are introduced in order to control the flow in and out of the chamber via the side channels (also labeled in red). The mechanism of the circular valves is the same as the “O-shaped” control valves for the payload channels, except there are no bypass channels in this layer. Instead, the circular valves were designed to fully block the flow in or out of the Culture Chamber if necessary.

Similar to the automated operation of the addressable ports, the circular control valves can be opened or closed on demand using the control software (see Section II.3). This allows the device to replenish the old media in the Culture Chamber without the need to actuate the addressable ports. Specifically, fresh media can be delivered to the Culture Chamber through the inlets (see green arrows in Figure 7A) at fixed intervals (e.g. every 5 hours) and old media can be flushed out through the outlets (see purple arrows in Figure 7B). Additionally, both control valves can be closed at the same time in order to prevent any fluid currents inside of the Culture Chamber. This is extremely important at the seeding stage, when the cells require a static condition in order to form focal adhesions with the substrate.

**Figure 7.**
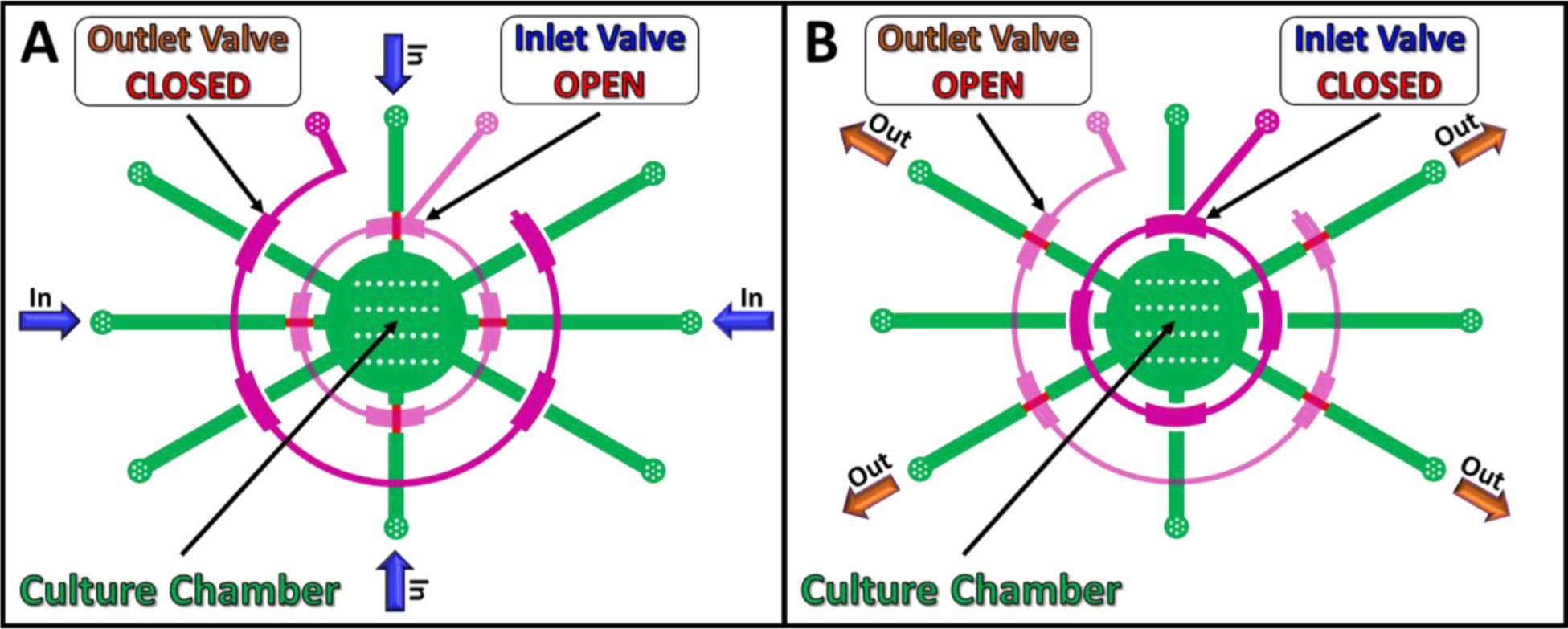
Top XY view schematic of the ‘Culture Chamber’ with the ‘Circular Control Valves’. The Culture Chamber is shown in red, and the Circular Control Valves surrounding it are in blue. **(A)** Fluid and cells can be added to the Culture Chamber through the 4 inlet channels (marked with green arrows) when the inlet Control Valve is “open” and the outlet Control Valve is “closed”. **(B)** Conversely, the 4 outlet channels (marked with purple arrows) can be used to remove fluids and cells from the Culture Chamber when the inlet Control Valve is “closed”, and the outlet Control Valve is “open”.

### III.2 Non-Disruptive *Fluid* Manipulation within the Culture Chamber

Some possible fluid and/or cell manipulations within the device are: 1) Seeding different cell types in varied amounts by flowing them into pre-determined spatial patterns; 2) Nourishing the cells in the Culture Chamber by continuously renewing the media; 3) Inducing and directing cell migration by establishing a dynamic nutrient and/or a chemoattractant gradient; 3) Patterning tissue by modulating cell differentiation and/or morphology via delivery of bio-agonists (e.g. growth, differentiation factors) and/or drugs (e.g., cytoskeleton-altering) to specified locations/selected cells within the device; and 4) Sampling a living culture non-disruptively by collecting and, sending off for analysis, effluents from different locations above the cells in the Culture Chamber.

The action of delivering and sampling chemicals within the addressable device is shown in the left and right panes of Figure 8, respectively. In the former case, a purple dye is delivered to the right bottom corner of a 4×4 array of microfluidic ports; while in the latter case, the same purple dye is withdrawn back via a port at the opposite end of the same address row. As a possible application, the picked-up fluid could be a cell culture effluent, which would then be sent off to an external sensor for ex-situ analysis. This would eliminate the reliance on destructive chemical assays (e.g., histological sectioning or crushing the sample for plate reader analysis) ensuring continuous monitoring of the biology occurring within live cultures. Furthermore, this can be done continuously over long periods of time, given that the whole process is automated (and as such, does not require any human involvement). Video 1 shows the operation of the device over time, where the purple dye is delivered and sampled to/at different addresses within the 4×4 grid of microfluidic ports.

**Figure 8.**
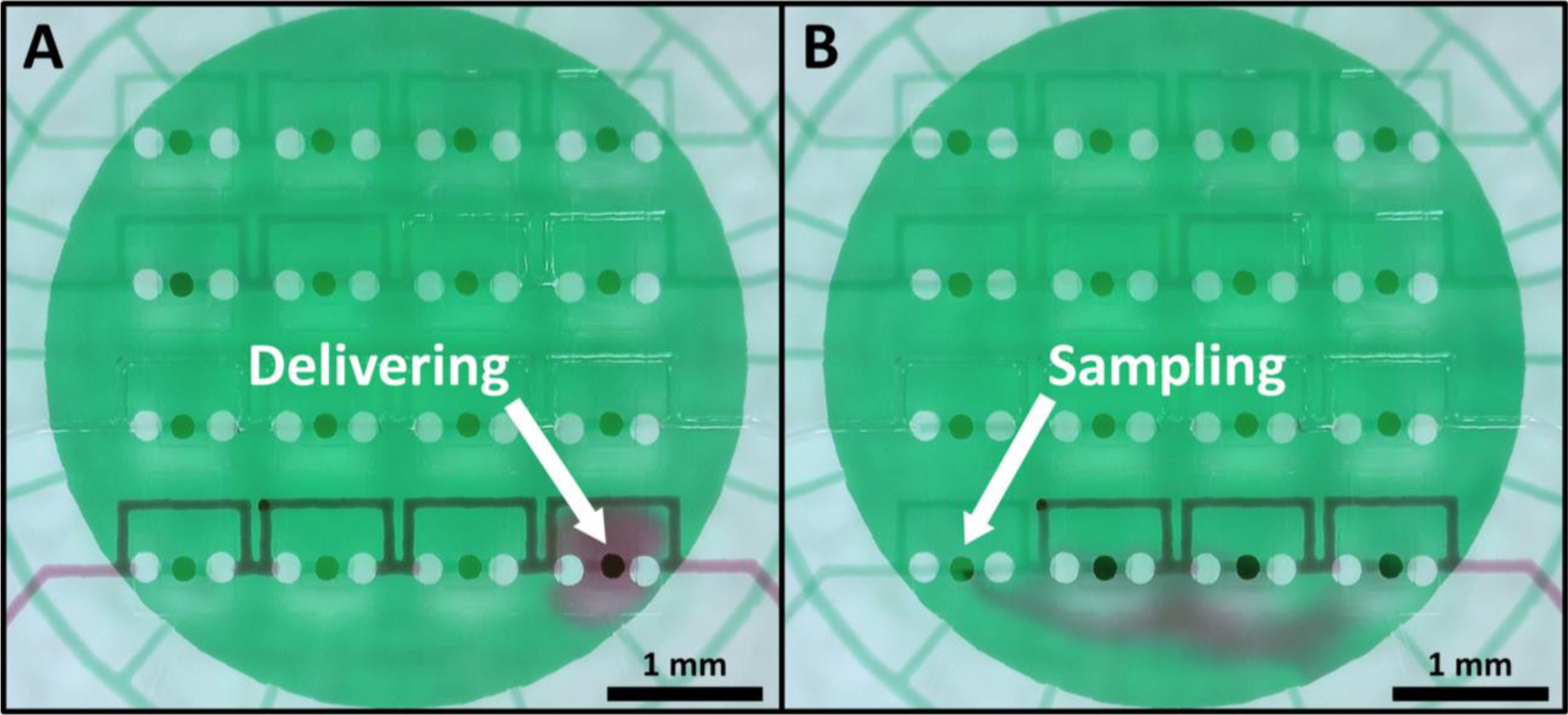
Microscopy of the fluid manipulation within the Culture Chamber. **(A)** Delivery of a purple dye to the bottom right corner port in the 4×4 grid; **(B)** Sampling of the delivered purple dye via the bottom left corner port in the 4×4 grid. The Culture Chamber is filled with green dye for contrast. Each port is 150 µm in diameter, and the channels are 100 µm in width. The view is from the top of the device.

### III.3 Minimally-Disruptive *Cell* Manipulations within the Culture Chamber

To further demonstrate the addressable device’s ability to manipulate *cells* within the Culture Chamber, its ports were used for seeding (i.e., additive manufacturing) a co-culture of mouse MSCs and NIH3T3 cells in a predetermined spatial pattern. Video 2 shows the process of the cell delivery through a single port to the Culture Chamber below, and Figure 9A shows the MSCs (green) seeded in a square shape that surrounds the NIH3T3s (red) deposited in its center. In this figure, each of the addressable ports in the 4×4 grid from Figure 6A was used to deliver the different cell types to the Culture Chamber below.

**Figure 9.**
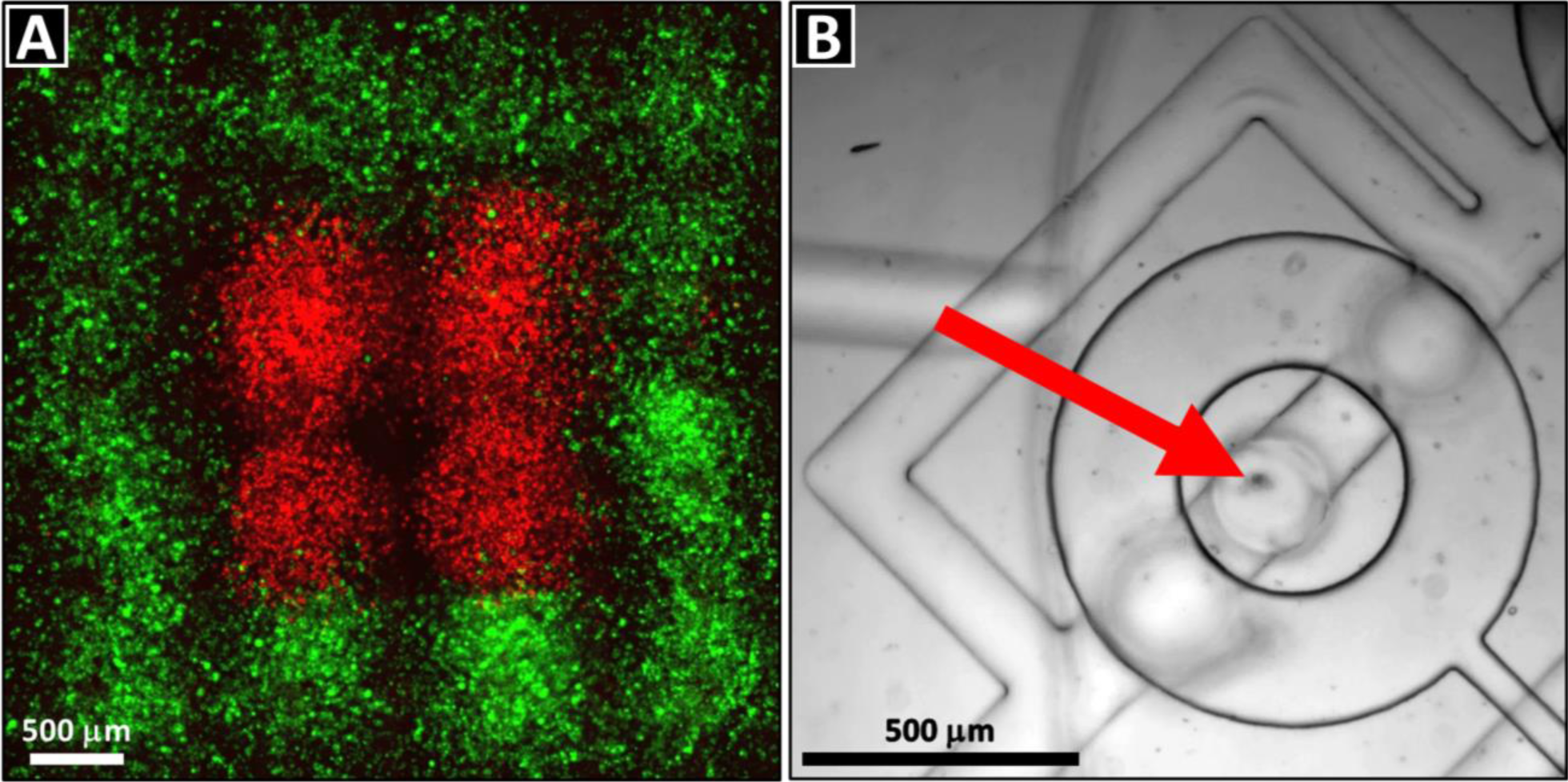
Minimally-disruptive additive cell manipulations within the Culture Chamber of the device. **(A)** Fluorescence panorama showing MSCs (green) seeded in a square pattern, with NIH-3T3s (red) in its center. Pane size is ∼4.5 x 4.5 mm. **(B)** Bright field microscopy image showing the ability to trap and manipulate a single cell (red arrow) at a time.

Although the resulting resolution is slightly lower than that of a typical bioprinter (e.g., 100-300 µm), it can be improved further by either reducing the feature size of the device’s design, or by controlling how many cells are being manipulated at a time. For example, Figure 9B and Video 3 show the device’s ability to trap single cells by the O-valves surrounding its microfluidic ports. This means that, in principle, the additive manufacturing can be done with a single cell precision (especially if machine vision-based automation is utilized). Alternatively, the seeding density can be controlled by varying the concentration of the cells in the carrier fluid and by changing how much of it can pass through an address while it is open. For example, Figure 10 shows a cell density gradient created by delivering MSCs to a row of addresses in progressively increasing amounts, from left to right. This was achieved by keeping the control valves open for progressively longer times during the cell delivery via flow: starting with 0.5 seconds for the left-most address, and then increasing the interval by 0.5 seconds for each subsequent location (ending with 2 seconds for the right-most one).

**Figure 10.**
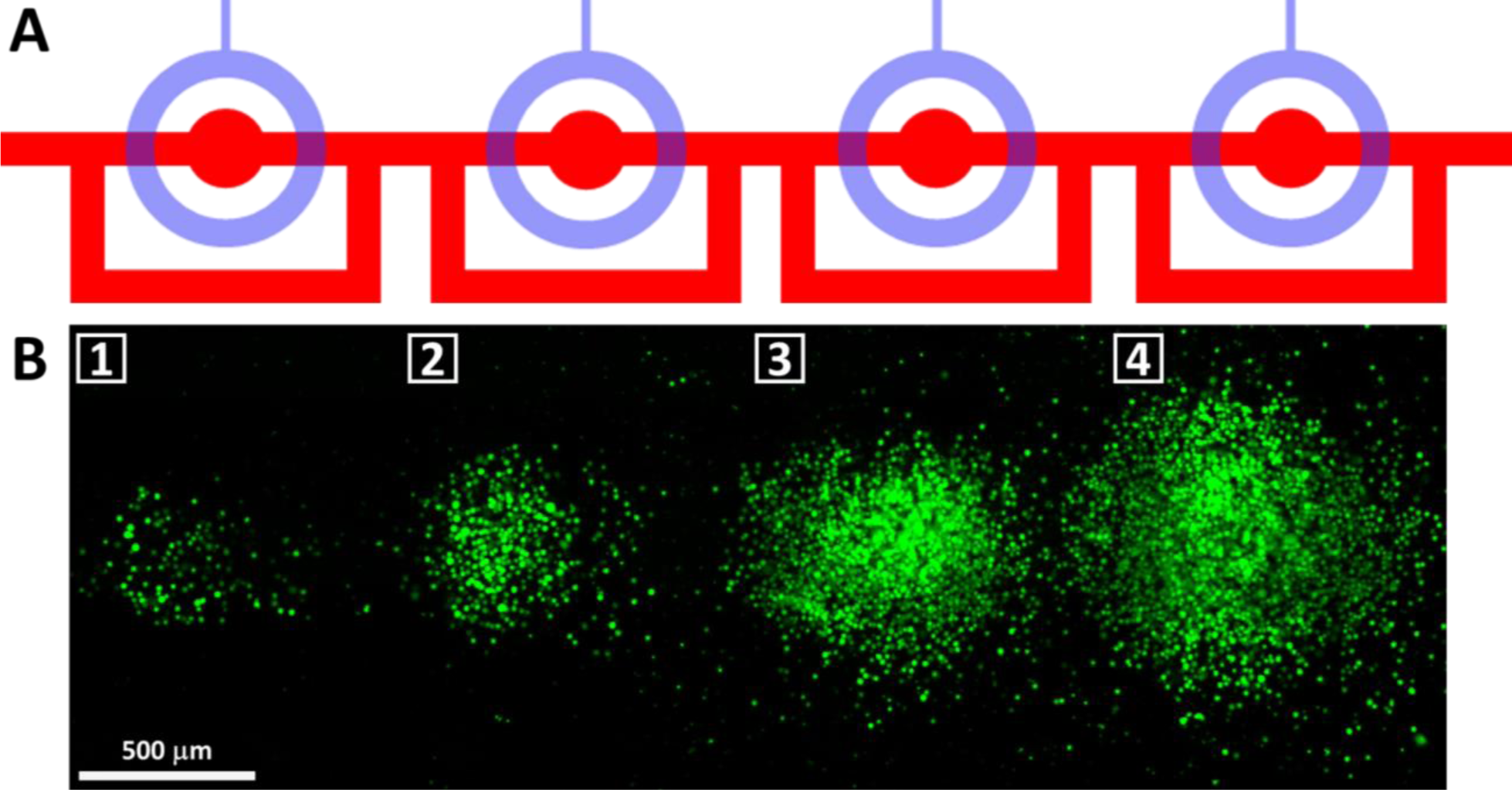
A cell seeding density gradient created by the device in order to demonstrate its utility in additive manufacturing. **(A)** Reference diagram showing the four addresses that were used for creating the cell seeding density gradient pattern. **(B)** Fluorescent microscopy of the MSC seeding gradient created in the Culture Chamber of the device, increasing in cell density from left to right. Panorama is ∼1.3 x 4 mm.

Additionally, Figure 11 demonstrates the device’s ability to create *inverse* cell patterns via *subtractive* manufacturing. The idea is to first seed cells uniformly in the Culture Chamber, and then let them adhere sufficiently to its floor in order to reach confluence (see Figure 11A). Subsequently, the focal adhesions anchoring the confluent cells to the bottom are cleaved by delivering a mixture of 0.25% Trypsin 0.2 g L^−1^ EDTA for a duration of 10 minutes. Eventually, the detached cells are flushed away via flow of PBS from user-specified addresses in order to create localized clearings arranged in a spatial pattern within the Culture Chamber in the culture below them (see Figure 11B). It should be noted that the PBS flow is is at times pulsed in order to increase the detachment of the cells from the substrate (see Video 4).

**Figure 11.**
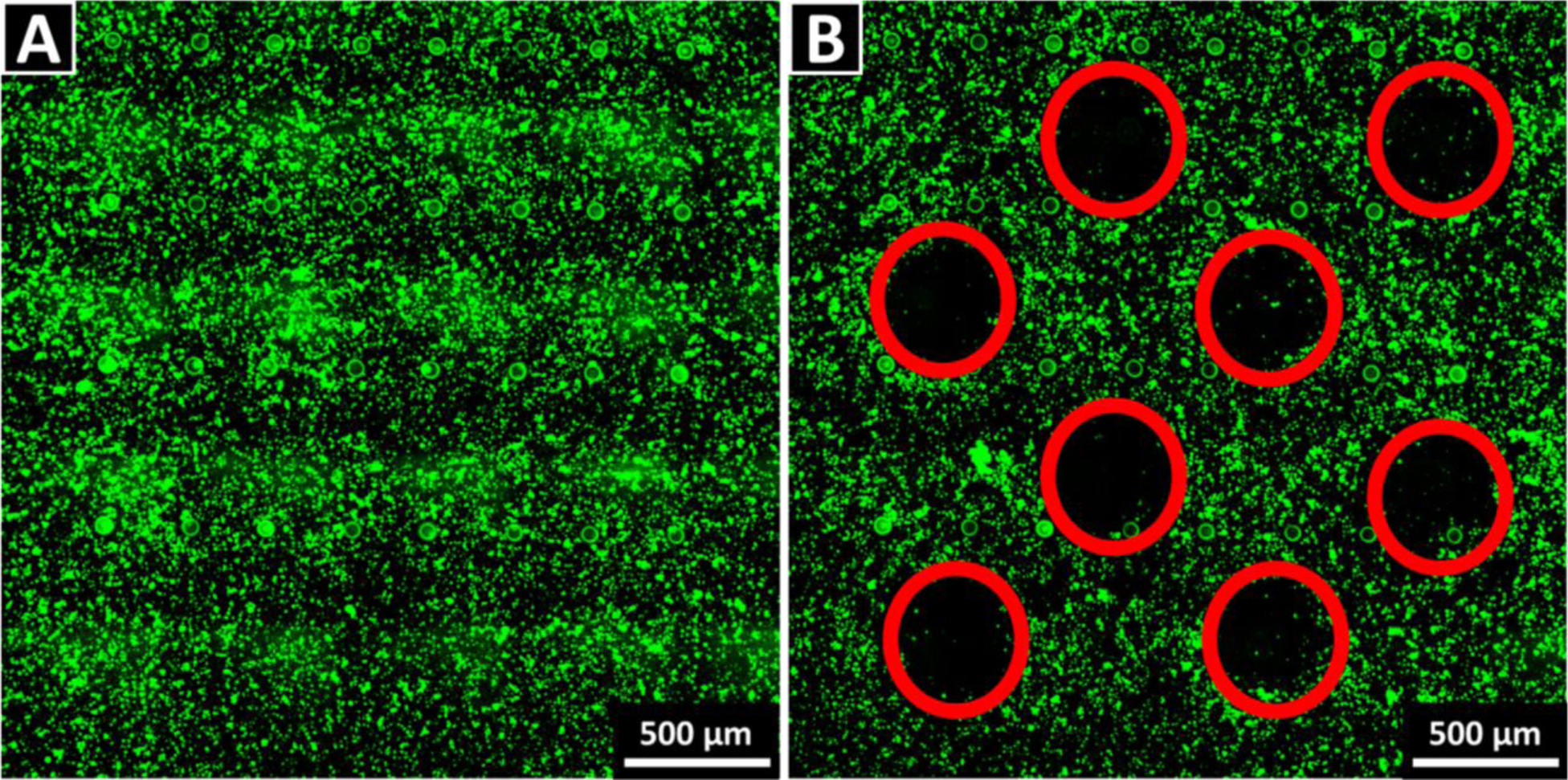
Fluorescent microscopy of minimally-disruptive subtractive manufacturing: **(A)** Culture Chamber’s “floor” covered with a confluent MSC layer (labelled using a fluorescent CellTracker™ Green CMFDA Dye) below a 4×4 matrix of addressable ports, before the cell subtraction; **(B)** Array of spaces (highlighted using red circles) from which the cells were removed via the subtraction (i.e., trypsinization followed by pulsatile flow). See Video 4 for an animation of the process at a representative addressable port.

This example of subtractive manufacturing demonstrates the ability of the device to remove undesired cell overgrowth and correct any localized seeding mistakes (something that is not possible with conventional scaffold and bioprinting approaches). Another possible application of this technology is the creation of contactless high throughput wound healing assays. Specifically, in a typical assay, it is desirable to be able to create an artificial “wound” opening in a monolayer of a patient’s cells grown to confluence. Subsequently, the cells surrounding the wound proliferate and migrate, eventually covering up the empty space and sealing the “injury”. Typically, these wounds are created using mechanical (e.g., scratch, stamp), thermal, electrical or optical damage to the cells.^36^ Most recently, flow focusing using microfluidics has provided a contact-free alternative for selectively removing cells enzymatically, in order to generate a wound with a clear boundary and without damaging the surrounding cells or the substrate’s surface coating (which is often necessary for increasing cell adhesion and/or directing the culture’s fate).^37–43^

However, all of these assays yield only linear-shaped wounds, whereas the addressable technology reported in this manuscript can create circular ones (as in Figure 11B). Finally, Video 5 shows the device’s ability to perform minimally-disruptive cell *biopsies* at different locations in the culturing chamber. This is inspired by how adherent single cells are trypsinized and then retrieved from cultures via micropipette suction.^44^ Similarly, in our device the adherent cells are trypsinized using the same procedure as in the subtractive manufacturing. However, in this case, they are drawn into the addressable port above via pressure reversal and are flowed away to a different location (e.g., for *ex-situ* analysis) via the microfluidic channels. Put together, these fluid and cell manipulation abilities demonstrate the versatility and the potential of our device to perform automated minimally-disruptive observations within living cultures.

## IV. DISCUSSION

The goal of this work was to create a technology capable of real time nondisruptive manipulation of cells and fluids within living cultures of artificial tissues. A major disadvantage of the existing biomanufacturing methods is the disconnect that currently exists between the scaffold fabrication and its culturing: namely, once the cells are either seeded or bioprinted into conventional scaffolds, all access to them is lost for the remainder of the culturing duration. As a result, the existing controls over cell behavior are typically done in one of the following ways: A) Scaffolds – substrate shape, stiffness, charge, porosity and chemical composition are used to affect cell fate and tissue development; B) Biologicals – various growth and differentiation factors, antiseptics and other drugs / biomolecular agonists are either added to the cell culture media, or released from the scaffold over time; the cells’ DNA is modified to express or knockout a certain feature of interest; foreign microorganisms are introduced to produce synergic interactions and signaling queues; and C) Physicomechanical – shaking, mechanical loads etc. are applied to enhance the cultures due to a stimulatory effect that they have on the cells.^45^

However, the overarching limitation of these *fabrication-based* controls is that they are either *static*, in the sense that they do not vary over time (e.g. substrate material, shape); or, are “blind” (aka, *open-loop*), as in the case of the timed-drug release from scaffolds. In other words, the stimuli do not adapt in response to the cell behavior. Furthermore, after the cells are either seeded or bioprinted into the scaffold, they cannot be targeted precisely. Consequently, only *bulk level* monitoring (e.g., probing the temperature and the pH of the bioreactor effluent) and controls (e.g., adjusting the media contents and the bioreactor environment) are available during the culturing stage of the artificial tissue manufacturing. So, the culture-based controls are not only poor in a sense that there is little feedback available, but they have negligible spatial resolution as well. Yet, precise closed-loop controls over *localized* cell behavior throughout the *entire* culturing process are necessary for producing viable tissue and maintaining product consistency. For example, in vivo organogenesis occurs in *multiple-steps*; this is illustrated by how most bones in our bodies start out as cartilage, and only subsequently become calcified through a process called endochondral ossification. Thus, in order to replicate such dynamic processes in vitro, both *spatial* and *temporal* control over the tissue development is required.

For this reason, several attempts have been made in the past to create the microfluidic scaffolds;^11–18^ most notably, Prof. Robert Langer’s,^12^ who is regarded as one of the founders of tissue engineering. The idea is that the integration of “vasculature” into the scaffolds can potentially resolve product size limitations and yield consistent tissue quality. Specifically, a combination of micro channels and ports could be utilized for patterning tissue by flowing different types of cells into the desired positions; overcoming product-size limitations^46^ by nourishing the cells deep within *large* scaffolds; enabling long-term survival of the cells in artificial environments by clearing metabolic waste from the scaffold pore space; and controlling the tissue growth in an adaptive manner by modulating the cell behavior, based on nondestructive microscopy and ex-situ chemical assay analysis (which could also be accomplished by using the microfluidic ports to sample the culture effluent). These abilities would result in an improved cell viability compared to conventional biomanufacturing methods (e.g., in bioprinting many cells die as a result of the need to participate in the fabrication process), and in much needed spatiotemporal control over the cell behavior within the living cultures. This would in turn clear major bottlenecks to the advancement of organ manufacturing and drug testing using organ mimics. However, the past work on the microfluidic scaffolds has focused only on developing *materials* for these devices,^11–, 18^ while the microfluidic plumbing necessary for the targeted nondisruptive fluid and cell manipulations at different locations within the artificial tissue constructs has not been designed.

To that end, we hypothesized that an *addressable* plumbing design, commonly used for high throughput microfluidic assays, could be modified in order to enable the minimally-disruptive cell and/or chemical delivery and/or removal/sampling, at targeted locations within the artificial tissue scaffolds. As proof of concept, we fabricated a prototype device, which can perform the said operations within a 2D living cell culture. Furthermore, we automated the device, in order to eliminate the reliance on human labor over the long-term culturing periods that are typically expected in tissue engineering experiments. Consequently, we have shown that the device is capable of a localized payload (both fluid and cell) delivery and *sampling*, directly below any desired address. Furthermore, we have used this technology to demonstrate both *additive* and *subtractive* manufacturing, by creating various co-culture seeding patterns and using trypsin delivery to detach the cells at selected locations. None of these operations are currently possible in tissue scaffolds fabricated using conventional means (e.g., de-cellularized organ, bioprinted, etc.), after the cells have been seeded into their pore spaces. Yet, the resolution with which the tissues could be created and patterned are already comparable to that of bioprinters. Moreover, it could be improved even further by creating a microfluidic scaffold with smaller plumbing features via advanced fabrication techniques; and, by introducing machine vision algorithms into the automation software, which would increase the cell trapping precision via the O-valve beyond what is currently practical manually.

In principle, single cell behavior such as migration, differentiation, proliferation and tissue deposition could all be controlled using this approach, throughout the entire culturing process, which is currently not possible using the existing biomanufacturing methods. Moreover, this complementary “marriage” of the two technologies would blur the line between fabrication and culturing stages of the tissue-manufacturing process. Instead, it would make the entire process *continuous*, thereby allowing to retain the precision, and the control over individual cells in the scaffold, well into the culturing stage. However, there are still technological difficulties that need to be overcome in order to advance the 2D prototype to a fully-fledged 3D scaffold: namely, 1) The addressable microfluidics plumbing needs to be scaled up for XYZ delivery / sampling (instead of just XY); 2) To facilitate mass production of the 3D scaffolds, a manufacturing method that is free of manual layer-by-layer alignment needs to be developed. There are several potential candidate technologies that could be used for this in the near future. Namely, digital micro-mirror device projection printing (DMD-PP) is a stereolithographic method that uses an array of micro-mirrors to spatially modulate a pulsed UV laser, in order to achieve additive crosslinking patterns in a photo-curable material.^16, 47–50^ Similarly, it can also be used for subtractive laser ablation within previously crosslinked layers of the material. While an alternative approach is to 3D print a *sacrificial* mold (a.k.a. “template”), shaped like the inverse of the desired scaffold pore network.^15^ The template is then submerged into a gel precursor of the scaffold material. After the gel has solidified around the template, the latter is dissolved out of the gel, leaving behind the desired porous networks in the scaffold. 3) Finally, the scaffold should be biocompatible and *biodegradable* when implanted in vivo. As mentioned before, there are a number of candidate materials that have been custom-designed specifically for the purpose of fabricating the microfluidic scaffolds: such as poly(ester amide), poly(1,3-diamino-2-hydroxypropane-co-polyol sebacate) (APS)^14^, poly(glycerol-co-sebacate) (PGS),[13] poly(octamethylene maleate (anhydride) citrate) (POMaC),^17^ Poly(ethylene glycol) diacrylate (PEGDA),^15^ poly(l-lactic-co-glycolic) (PLGA)^11^, and Silk fibroin^13, 18^. However, given that the scaffolds in these studies did not utilize the use of microfluidic valves, it is not apparent whether these materials are stiff enough to make them. Only the silk fibroin has been used to create such valves, but it is not a cross-linkable material,^18^ so it is currently not compatible with the 3D stereolithographic fabrication methods such as the DMD-PP. Therefore, further work should be done in order to develop new cross-linkable, biodegradable/biocompatible, materials that are stiff enough to create the microfluidic valves, and that would ideally be optically transparent for microscopy observation.

Overall, the successful development of the microfluidic scaffolds will help to minimize the crippling tissue variability plaguing the biomanufacturing industry today. This will benefit patients in need of life-saving transplants by accelerating the translation of regenerative medicine technologies to the clinical market. Moreover, the minimally-disruptive visual and chemical observation of the cell behavior in the 3D artificial tissue cultures will lower experiment costs and provide data-gathering continuity superior to the conventional analysis (i.e., destructive assays for each time point). Ultimately, the proposed approach could pave the way toward tracking and controlling of every cell in a culture on an individual basis, given that its targeting precision is limited only by the fabrication quality of the scaffold (to be refined tremendously by the anticipated explosion of 3D printed microfluidics[6, 7]). We expect that this will shift the tissue engineering paradigm from integrating “blind” (i.e., open-loop) controls into the scaffold’s material (e.g., timed drug release) to controlling cells in growing tissues interactively instead. Finally, the computer-driven culturing would solve a major logistical hurdle to the adoption of such technologies by allowing companies to provide digital codes to hospital staff, instead of having to teach them custom culturing protocols for each new product (which is impractical). Therefore, this technology would both simplify the process for the end-user and ensure specification-compliance on-site.

## V. CONCLUSION

We have presented a microfluidic platform capable of performing spatiotemporal manipulations of fluids and cells within a living culture. Specifically, we have interlaced a PDMS prototype with addressable microfluidic ports and demonstrated that it is capable of delivering / sampling fluids in the Culture Chamber situated at the bottom of the chip. Furthermore, we have shown *additive* manufacturing by creating spatial cell seeding patterns by flowing fluorescent MSCs and NIH3T3s into different locations within the device. Moreover, we also demonstrated *subtractive* manufacturing by detaching selected cells via localized trypsin delivery and flushing them from the Culture Chamber. Lastly, we showed that the detached cells can also be captured and sent off ex-situ, thereby overcoming the need for destructive analysis that is common in tissue engineering studies. In the near future, we will extend this concept to XYZ manipulations within transparent, biocompatible and biodegradable 3D scaffolds. It is our hope that the design will ultimately resolve the bottlenecks plaguing the conventional biomanufacturing technologies today, and ultimately enable computer-driven tissue engineering through coupling with automation electronics. Specifically, the proposed technology would enable: 1) organ-sized engineered tissue via nutrient delivery and metabolic waste removal throughout the scaffold; 2) viable and consistent tissue products via orchestration of cell action during culturing; 3) improved drug screening via nondestructive sampling of cell responses throughout the entire time course of administration; 4) biomanufacturing automation for achieving desired specifications on site, without the need to train hospital staff custom culturing protocols for every different tissue engineering product. Overcoming these bottlenecks would in turn allow machines to culture complex artificial organs viably and reproducibly, which is the ultimate goal of tissue engineering.

## Supporting information

Video 1 - Sampling and Deliverying of Dyes

Video 2 - Close Up of Cell Delivery Through an Address

Video3 - Single Cell Manipulation

Video 4 - Subtractive Manufacturing

Video 5 - Cell Biopsy

## ACKNOWLEDGEMENTS

The authors would like to thank Gustavus and Louise Pfeiffer Research Foundation Major Investment Grant, New Jersey Health Foundation Research Award - Grant #PC 22-19, NJIT Faculty Seed Grant, NSF I-Corps Site - Grant #1450182 for their gracious funding of our work. Additionally, the authors would like to thank NSF XSEDE EMPOWER for funding undergraduate labor, New Jersey Institute of Technology (NJIT)’s McNair Achievement, and Provost Summer Research Programs for providing student labor for this project. Finally, we would like to thank Dr. Rafael Gómez-Sjöberg for providing us with software and blueprints for the automated pneumatic pumping technology. We also acknowledge that computational resources were provided by the New Jersey Institute of Technology’s Academic and Research Computing Systems (ARCS) and Texas Advanced Computing Center (TACC) at the University of Texas at Austin through the Extreme Science and Engineering Discovery Environment (XSEDE)^51^ program, which is supported by National Science Foundation grant number ACI-1548562.

## VI. COMPLIANCE WITH ETHICAL STANDARDS

### Funding

This work was supported by the Gustavus and Louise Pfeiffer Research Foundation’s Major Investment Grant, New Jersey Health Foundation Research Award - Grant #PC 22-19, NJIT Faculty Seed Grant, and NSF I-Corps Site - Grant #1450182. Undergraduate labor was supported by the NSF XSEDE EMPOWER Program.

## Ethical approval

This article does not contain any studies with human participants or animals performed by any of the authors.

